# Whole genome methylation analysis of non-dysplastic Barretts oesophagus that progresses to invasive cancer

**DOI:** 10.1101/114298

**Authors:** MP Dilworth, T Nieto, JD Stockton, C Whalley, L Tee, JD James, MT Hallissey, R Hejmadi, N Trugdill, O Tucker, AD Beggs

## Abstract

**Objective:** To investigate differences in methylation between patients with non-dysplastic Barretts’ oesophagus who progress to invasive adenocarcinoma and those that do not.

**Design:** A whole genome methylation interrogation using the Illumina HumanMethylation 450 array of patients with non-dysplastic Barrett’s Oesophagus who either develop adenocarcinoma or remain static, with validation of findings by bisulfite pyrosequencing

**Results:** In total, 12 patients with “progressive” vs. 12 with “non-progressive” non-dysplastic Barrett’s oesophagus were analysed via methylation array. Fourty-four methylation markers were identified that may be able to discriminate between non-dysplastic Barrett’s Oesophagus that either progress to adenocarcinoma or remain static. Hypomethylation of the recently identified tumour supressor *OR3A4* (probe cg09890332) validated in a separate cohort of samples (median methylation in progressors = 67.8% vs. 96.7% in non-progressors,p=0.0001, z = 3.85, Wilcoxon rank sum test) and was associated with the progression to adenocarcinoma. There were no differences in copy number between the two groups, but a global trend towards hypomethylation in the progressor group was observed.

**Conclusion:** Hypomethylation of *OR3A4* has the ability to risk stratify the patient with non-dysplastic Barrett’s Oesophagus and may form the basis of a future surveillance program.

## INTRODUCTION

Oesophageal adenocarcinoma (OADC) incidence is increasing [1] and currently forms 5% of the digestive tract cancers in the UK [2]. Overall disease survival is poor [3] but correlates with stage of cancer at presentation, demonstrating significant survival advantages with detection of early stage disease [4, 5].

Barrett’s Oesophagus (BO), in which normal squamous mucosa changes to a columnar epithelium, results from prolonged exposure to stomach acids and bile salts which reflux into the oesophagus causing chronic inflammation and tissue damage [6].

The incidence of BO is increasing, largely thought to be a consequence of obesity induced reflux disease [7, 8, 9, 10]. Over time, driven by chronic inflammation, metaplastic change is observed and is associated with an increased risk of OADC [11]. For the majority of patients, BO will never progress beyond this benign metaplastic cell change [12, 13]. However, in a small number of patients, dysplasia will develop with some progressing to OADC [14]. The incidence of OADC in the BO population is up to 150 times greater than unaffected individuals [12].

Although the pathological changes seen in Barrett’s adenocarcinoma are understood as part of a well-established metaplasia-dysplasia-carcinoma sequence, a process driven by chronic inflammation [15], the molecular drivers are less clear The current dilemma is that for patients with non-dysplastic BO, there are no accurate methods for identifying the small number of patients at high risk of progression to cancer.

Surveillance practice of the Barrett’s patient varies widely, between some who endoscope patients each year in contrast to others who will never repeat the investigation[16]. The on-going MRC funded BOSS study aims to understanding the optimum surveillance strategy, randomising between prospective monitoring BO patients with frequent endoscopic assessment or a “watch and wait” policy [17].

Clearly, there is need for a method of risk stratification in these patients in order to facilitate a streamlined surveillance programme by identifying high risk non-dysplastic BO patients. Attempts at biomarker development for stratification of high risk Barretts oesophagus have focused on mutational change, specifically around the role of *TP53* mutation in predicting “high risk” disease [18] given its role as a driver in oesophageal cancer. However Fitzgerald et al [19] have convincingly demonstrated the presence of pathogenic *TP53* mutations in apparently normal squamous oesophageal mucosa thus making its role in progression to invasive adenocarcinoma unclear. However, the role of epigenetic change in the pathogenesis of Barretts oesophagus and oesophageal cancer is less well understood, but may well happen much earlier in the cancer developmental pathway and, as a direct result, provide a more appropriate target for both predicting its development and potentially arresting tumourgenesis, should a suitable epigenetic modulator be identified.

Multiple methylation markers have been identified which can discriminate between high risk and low risk BO including *APC/ p16* [20], *MGMT* [21], *PKP-1* [22], *TIMP3/TERT* [23], *RUNX3/HPP1* [24] [25] and *AKAP12* [26]. Agarwal *et al* [27] performed an MeCIP array based approach to compare the methylomes of progressor vs. non-progressing patients. In patients that progressed to invasive adenocarcinoma, their original biopsies began either with no dysplasia, indefinite dysplasia or low grade dysplasia, making comparison difficult. However, subsequent analysis of the top 25 differential methylation patterns found three gene regions with hypermethylation amongst the progression group (*Pro_MMD2, Pro_ZNF358* and *Intra_F10)* with a trend otherwise towards global hypomethylation, in keeping with other epithelial pre-malignant conditions [28].

While the studies reviewed do show variation in methylation between progressive vs. non-progressive BO, the methodology has been heterogeneous and few have conducted the study with a group of the same patients tracked over time.

## AIMS

To determine whether there are differences in methylation in patients between high risk non-dysplastic Barrett’s Oesophagus which will progress to cancer vs low risk Barrett’s

## METHODS

### Patients and Samples

Two sample cohorts were identified containing patients who either progressed to OADC from non-dysplastic BO or remained with non-dysplastic BO (NDBO) identified from a prospectively maintain database of patients with BO at Sandwell & West Birmingham NHS Trust. Inclusion criteria for the study were progressing patients with non-dysplastic Barretts oesophagus who, when observed over the study period, developed OADC. Samples were only included where there was a NDBO biopsy and then histology evidence that the patient developed adenocarcinoma. Non-progressing patients were identified from a biopsy of NDBO which when followed over time never progressed beyond NDBO. To be included in this group, the patient must have been in a surveillance programme for a minimum of 15 years and have serial biopsies over that period. They were also required to have still been alive and to have had a NDBO biopsy within 2 years of this study.

Patients were excluded from the study if BO material was only available as part of tumour associated BO or if dysplasia was identified in any BO biopsies.

All tissue used was formalin fixed paraffin embedded (FFPE) samples obtained from pathology libraries and prepared by the University of Birmingham Human Biomaterials Resource Centre (ethical approval 09/H1010/75). H&E stained slides were reviewed by a consultant pathologist to ensure that the samples were BO and had no dysplasia throughout their extent. Cut 5μM paraffin sections were mounted onto frosted slides, and macrodissection for BO carried out. DNA extraction was then performed using Qiagen DNeasy Blood and Tissue kit following the manufacturer’s protocol. Each sample of extracted DNA was then quantified and qualified by Nanodrop spectrophotometry and Qubit fluorimetry. Bisulphite conversion was performed using a Zymo DNA Methylation bisulphite conversion kit following the modified Illumina Infinium protocol on 500ng of extracted DNA.

### Methylation Arrays

The Illumina HumanMethylation 450 array, in which the methylation status of more than 485,000 individual CpG sites are examined [29] was used to compare sample groups (progressing NDBO vs non-progressing NDBO). Once bisulphite converted, 1ng of DNA was quality controlled (Illumina FFPE QC kit) with only samples with dCt <5 being taken forward to array analysis. The resulting samples underwent repair suitable for array hybridisation using the Illumina FPPE restore kit. [30] followed by hybridisation to Illumina HumanMethylation450 arrays using manufacturers protocols and scanned on an Illumina iScan. Normalised intensity files (iDAT) were exported using GenomeStudio for downstream analysis.

### Immunohistochemistry

In order to understand the effect of methylation on the expression of OR3A4 correlation of OR3A4 expression as determined by immunohistochemistry (IHC) vs. progressor/non-progressor status was carried out. IHC was carried out on a Leica Bond RX system using the a mouse polyclonal anti-OR3A4 antibody (Abcam ab67107) at a dilution of 1:100 with a primary incubation time of 15 minutes.

IHC was scored on epithelial and stromal components and a composite score consisting of the sum of expression within membranous, nuclear and cytoplasmic compartments on a score of 1-4 was made. Scoring was carried out by two independent observers blinded to progressor/non-progressor status.

### Bioinformatic Analysis of Array Data

Bioinformatics analysis of the methylation microarrays was carried using the ChAMP package [31] via Bioconductor/R. In brief, red/green intensity values were captured from Illumina iDAT files, background corrected and SWAN normalised to produce M values.

M values were analysed using a logistical regression model using Empirical Bayesian shrinkage of moderated t-statistics to correct for small sample size. Small sample size was controlled for by setting stringent FDR Q-values of <0.05. Identification of variable methylated sites allowed the CpG site markers to be highlighted. DNA Copy number analysis was carried out using the *DNAcopy* module of ChAMP.

### Pyrosequencing validation of hits

Methylation insensitive primers were designed and sourced,using Qiagen PyroMark Primer Design software v2.0. Primers were designed to flank CpG sites of interest. Illumina CG methylation probe locations were retrieved from the UCSC genome browser [32] and FASTA sequence retrieved for -200bp to + 200bp of the target CG dinucleotide. Primer design settings were optimised to design amplicons suitable for FFPE pyrosequencing, with the optimum amplicon size set to between 80-150bp. All other settings were as per the standard Qiagen design parameters. Primers were ordered from Sigma Aldrich, with the biotinylated pyrosequencing primer being purified by high performance liquid chromatography and the remainder by desalting. Pyrosequencing PCR was performed using Qiagen PyroMark PCR Gold kit, consisting of 2uL of bisulphite converted DNA, 25uL of PCR master mix, 5uL of CoralLoad dye, 3uL MgSO_4_, 10uL of Q reagent and 2.5uL each of forward (20mM) and reverse (20mM) primer. Reaction conditions were determined experimentally by use of a gradient PCR for each primer pair. A typical reaction consisted of activation at 95C for 15 minutes, followed by 45 cycles of denaturation at 94C for 30s, annealing at 56C for 30s and extension at 72C for 30 seconds, followed by a final extension step at 72C for 10 minutes. In addition to experimental DNA, each PCR was performed with 100% methylated DNA, 100% un-methylated DNA and ddH2O as controls. Methylated and unmethylated DNA was generated in house by means of M.SSl conversion (methylated DNA) and whole genome amplification using the Qiagen Repli-G kit (unmethylated DNA). Primer sequences for validation pyrosequencing were as follows. For FGFR2 cg17337672 these were Foward = AGGGGAAGGGAATTTAGGTT; Reverse =

[Btn]TCAATCTTCCCCCAAACAACCACT and sequencing =

GTTTAGAAGTTTTTTTTGGATTAGT. For ORA3A4 cg07863524 these were forward

= GTGGTAGAAGTAGGATGAGGTGTTGATAAT; Reverse

=[Btn]CTTCAACTTCCTTCCCCTTACATTT and sequencing =

GGGTAGGGATGGAAGA. For OR3A4 cg09890332 these were Forward =

TTAAAGTGTTAGGATTATAGGTGTGAGTTA, reverse =

[Btn]TTTCCCAACCCTAATCACTACTAATAAAAT and sequencing =

GGATTATAGGTGTGAGTTAT.

## RESULTS

### Patient selection

In total, 67 patients were recruited, 37 from Sandwell and West Birmingham NHS Trust (SWBNT) and 30 from University Hospital Birmingham NHS Trust (UHBFT). Of these, 20/67 progressed from non-dysplastic Barrettts oesophagus and 47 did not. The age range was between 42-60 years with a median age of 56 years of which 60/67 (89.6%) were of male gender. The median time to diagnosis of OADC in “progressive” patients was 114 months, with a range of 4-162 months. Of the patients recruited, in the SWBNT group 6 progressors and 6 non-progressors and in the UHBFT 6 progressors and 4 non-progressors were taken forward to methylation array analysis, giving a total of 12 progressors and 10 non-progressors. All patients recruited had validation pyrosequencing performed.

### Methylation microarray analysis

Twenty four samples in total were hybridised successfully to Illumina HumanMethylation450 microarrays. All arrays passed manufacturers QC as specified by metrics in Illumina GenomeStudio. Differential methylation analysis at the probe level (Table 1) revealed significant differences in methylation between progressor and non-progressors in non-dysplastic BO (Figure 1). In total, 44 significantly (defined as Bayes Factor, BF>5) differentially methylated targets were identified, the bulk being hypomethylated, with a trend towards global hypomethylation in progressor samples.

**Table 1.**
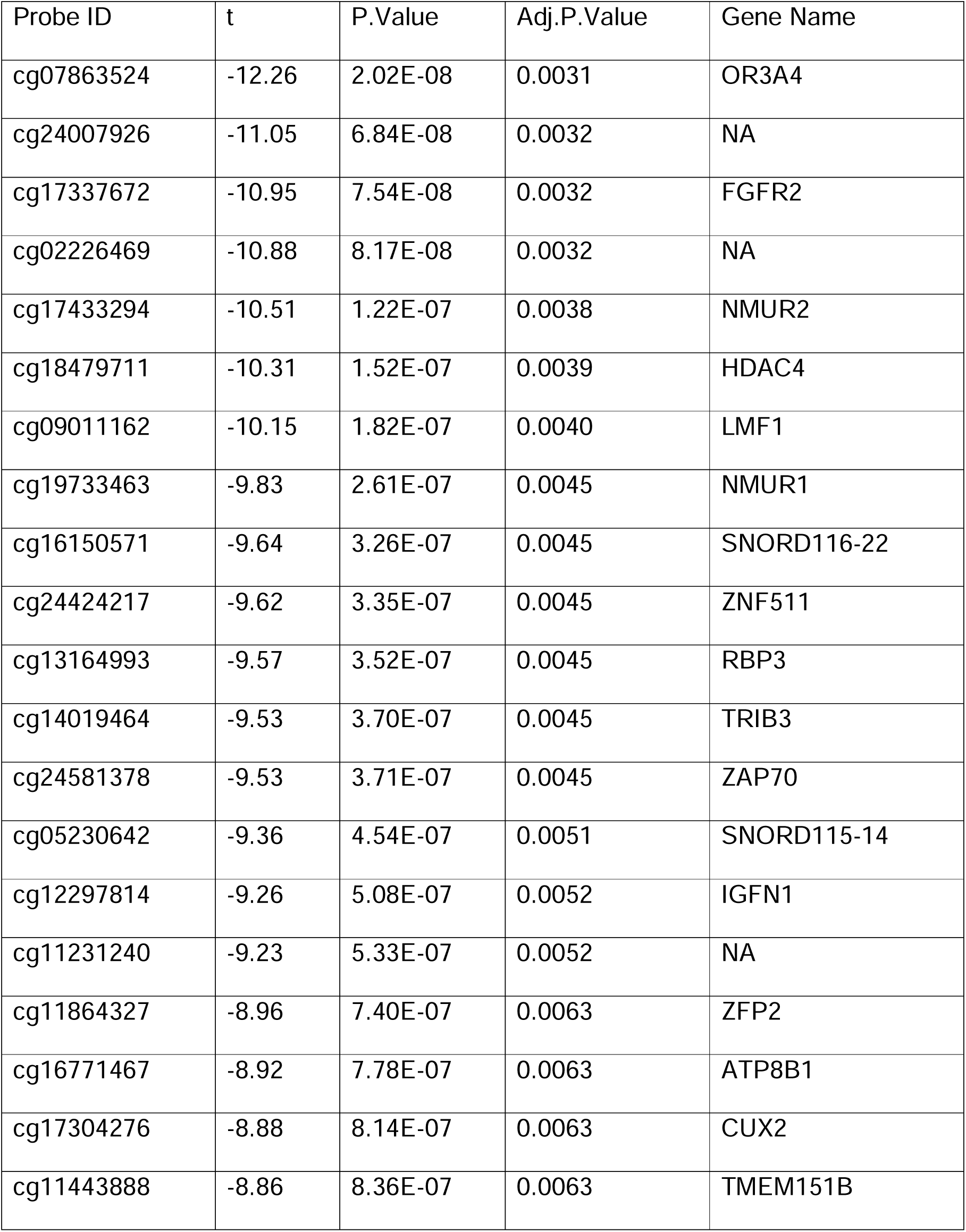
Top20 array identified CpG sites with methylation variation between non-dysplastic samples which progress to OADC vs those which remain static Key: Probe ID - the Illumina cg probe ID from the Illumina manifest, T = the t value - the size of the difference relative to the variation in the sample, p-value = the raw p-value, not corrected for multiple testing, Adjusted p-value = the p-value corrected for multiple testing

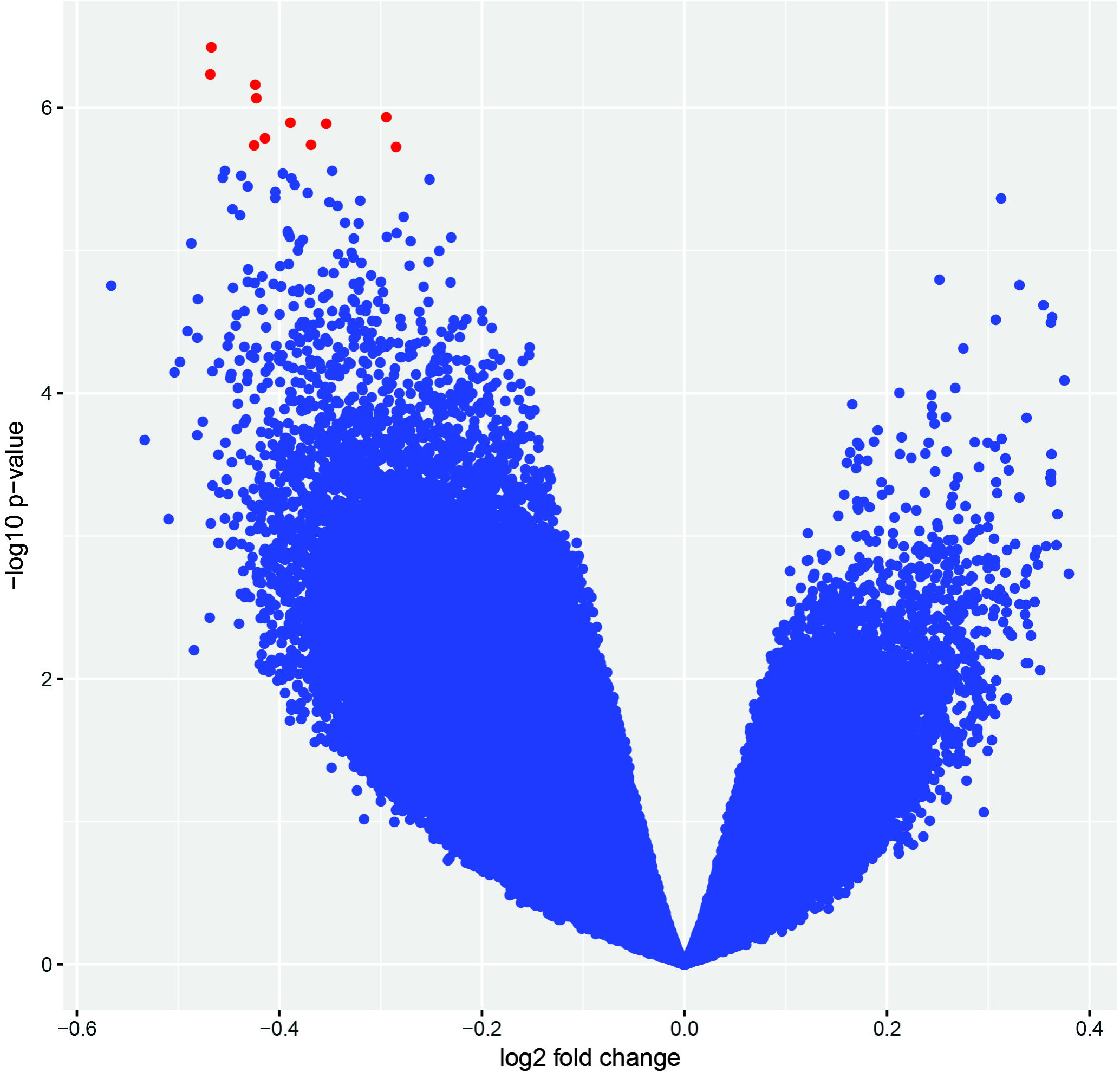

#### Differential methylation at the probe level

The top ranked differentially methylated probe was cg09890332 (chr17:3212495-3212495, hg19 coordinates) which tags a CpG dinucleotide -1044bp upstream of the TSS of the long non-coding RNA, *OR3A4* (NRR_024128.1). The second highest ranked differentially methylated probe was cg24007926 (chr2:206842761-206842761, hg19 coordinates. This CpG dinucleotide is with a large, intragenic region, with the nearest gene being *INO80D* (INO80 complex subunit D, NM_017759.4), 15,684bp downstream of this CpG. The third highest ranked differentially methylated probe was cg17337672 (chr10:123354172-123354172) which tags a CpG dinucleotide within intron 2 of *FGFR2* (Fibroblast growth factor receptor 2, NM_000141.4).

Differentially methylated regions (DMR) were called between progressors and non-progressors via the *dmrLasso* function of the CHAMP software package (Table 2). Significant DMRs were found from chr2:503065-503193 (which tags an intragenic region, DMR p= 7.69×10-4), chr5:8217236-8217322 (which also tags an intragenic region, DMR p=1.27×10-3) and chr10: 123353418-123355576 which spans a region from the 5’ UTR of *FGFR2* to the 1^st^ exon within *FGFR2* (DMR p=4.79×10^−3^).

**Table 2.** Table of differentially methylated regions (DMR)

| DMR ID | Probe ID | Probe level<br>adjusted p-value | Chromosome | Gene | Start of<br>DMR (bp) | End of<br>DMR (bp) | Size<br>in bp | Change in<br>methylation | P-value<br>for<br>DMR |
| --- | --- | --- | --- | --- | --- | --- | --- | --- | --- |
| 1 | cg21273584 | 0.014 | 2 | NA | 502999 | 503195 | 197 | -34% | 7.69E-04 |
| 1 | cg00854591 | 0.045 | 2 | NA | 502999 | 503195 | 197 | -16% | 7.69E-04 |
| 1 | cg11573608 | 0.014 | 2 | NA | 502999 | 503195 | 197 | -27% | 7.69E-04 |
| 2 | cg25568703 | 0.047 | 5 | NA | 8216903 | 8217655 | 753 | -25% | 1.27E-03 |
| 2 | cg25016964 | 0.039 | 5 | NA | 8216903 | 8217655 | 753 | -22% | 1.27E-03 |
| 2 | cg17642708 | 0.021 | 5 | NA | 8216903 | 8217655 | 753 | -33% | 1.27E-03 |
| 3 | cg10788901 | 0.172 | 10 | FGFR2 | 123352704 | 123355661 | 2958 | -27% | 4.79E-03 |
| 3 | cg14856220 | 0.044 | 10 | FGFR2 | 123352704 | 123355661 | 2958 | -24% | 4.79E-03 |
| 3 | cg17337672 | 0.003 | 10 | FGFR2 | 123352704 | 123355661 | 2958 | -47% | 4.79E-03 |
| 3 | cg02412684 | 0.031 | 10 | FGFR2 | 123352704 | 123355661 | 2958 | -38% | 4.79E-03 |
| 3 | cg06791446 | 0.058 | 10 | FGFR2 | 123352704 | 123355661 | 2958 | -13% | 4.79E-03 |
| 3 | cg22633036 | 0.581 | 10 | FGFR2 | 123352704 | 123355661 | 2958 | -4% | 4.79E-03 |

We took advantage of the information provided by the two colour Illumina Infinium chemistry to call copy number aberrations (CAN) within the regions targeted by the methylation probes using the CNA calling function of CHAMP. This did not demonstrate any recurrent copy number alterations between progressors and non-progressors. There were no significant differences in the numbers of CNA between the two groups, with a median of 43 CNA (range 24-92) in the progressors vs. 44 CAN (range 30-52) in the non-progressors (p=1.0, Wilcoxon rank sum).

Pathway methylation analysis was carried out using DAVID. Initially KEGG pathway analysis showed that genes associated with MAPK signalling were enriched in the dataset (p=0.012). Gene ontology analysis using the UP_KEYWORDS feature showed significant enrichment for the disease mutation (p=9.6×10-6), polymorphism (p=1.7×10-5), glycoprotein (p=3.4×10-5), and alternate splicing (p=1.1×10-4) terms.

### Validation pyrosequencing

Because of the likely biological relevance of *FGFR2,* and the data demonstrating that *OR3A4* was the top differentially methylated CpG, validation pyrosequencing was carried out on all 67 patients. Normality of distribution of methylation values was ascertained by histogram plots, in which it was found that methylation was non-normally distributed, therefore non-parametric testing was carried out For OR3A4 cg09890332, median methylation was 67.8% in progressors vs. 96.7% in non-progressors (p=0.0001, z = 5.158, Wilcoxon rank sum test) (Figure 2). The pyrosequencing assay design used covered two additional CpG +4bp and +10bp downstream of cg09890332. Median methylation in these were was 66.8% and 59.7% in progressors vs. 75.0% and 68.1% in non-progressors (p=0.0280 and 0.0368, z=2.197 and 2.088, Wilcoxon rank sum test). In order to investigate whether this phenomenon was localised to this region or was a gene-wide phenomenon, an additional pyrosequencing assay was designed based on probe ID cg07863524 (chr17:3213471-3213471) which is +976bp downstream from cg09890332 and -68bp from the TSS of OR4A4. This demonstrated that median methylation was 62.2% in progressors and 56.7% in non-progressors (p=0.600, z=−0.524, Wilcoxon rank sum test).

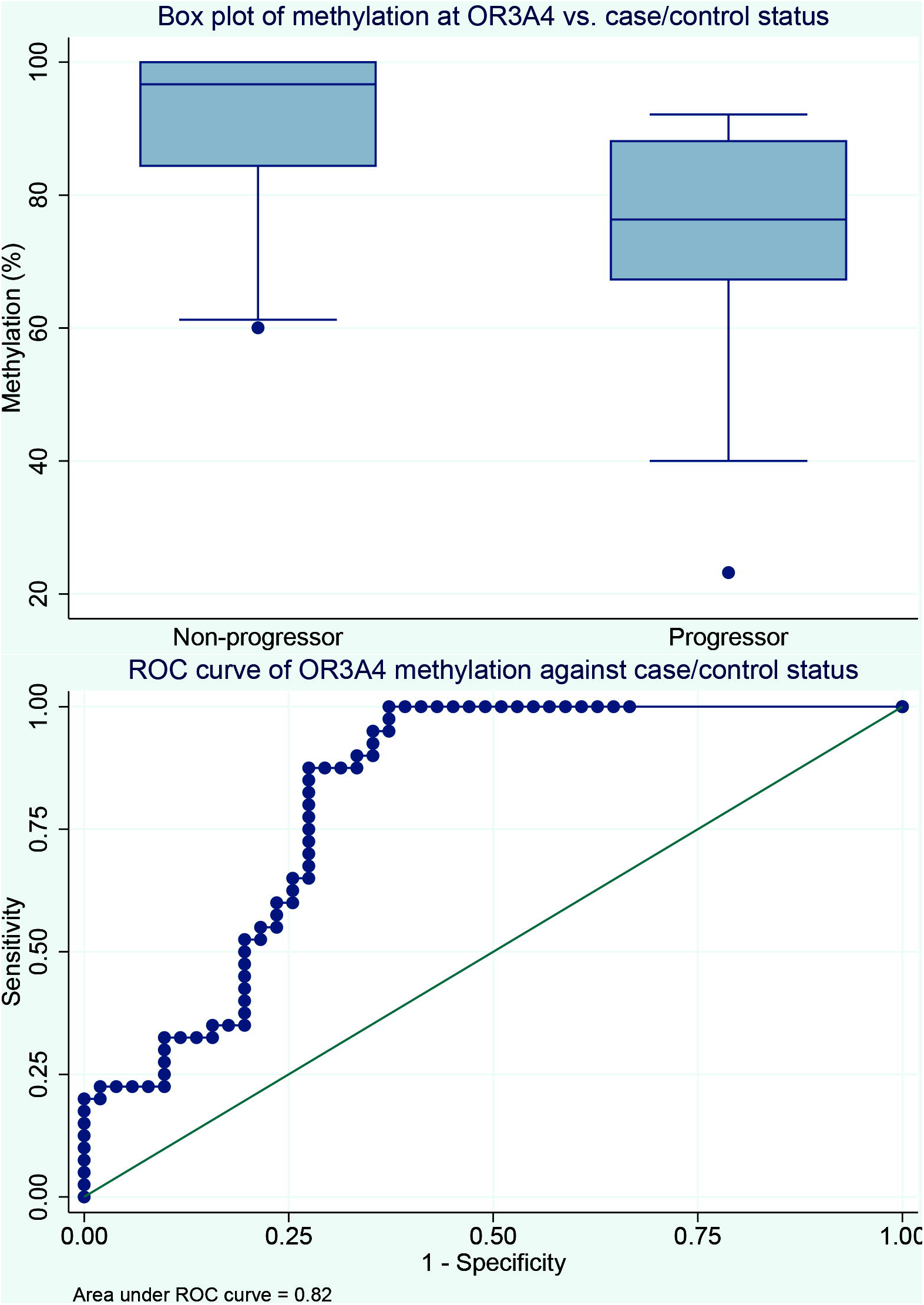

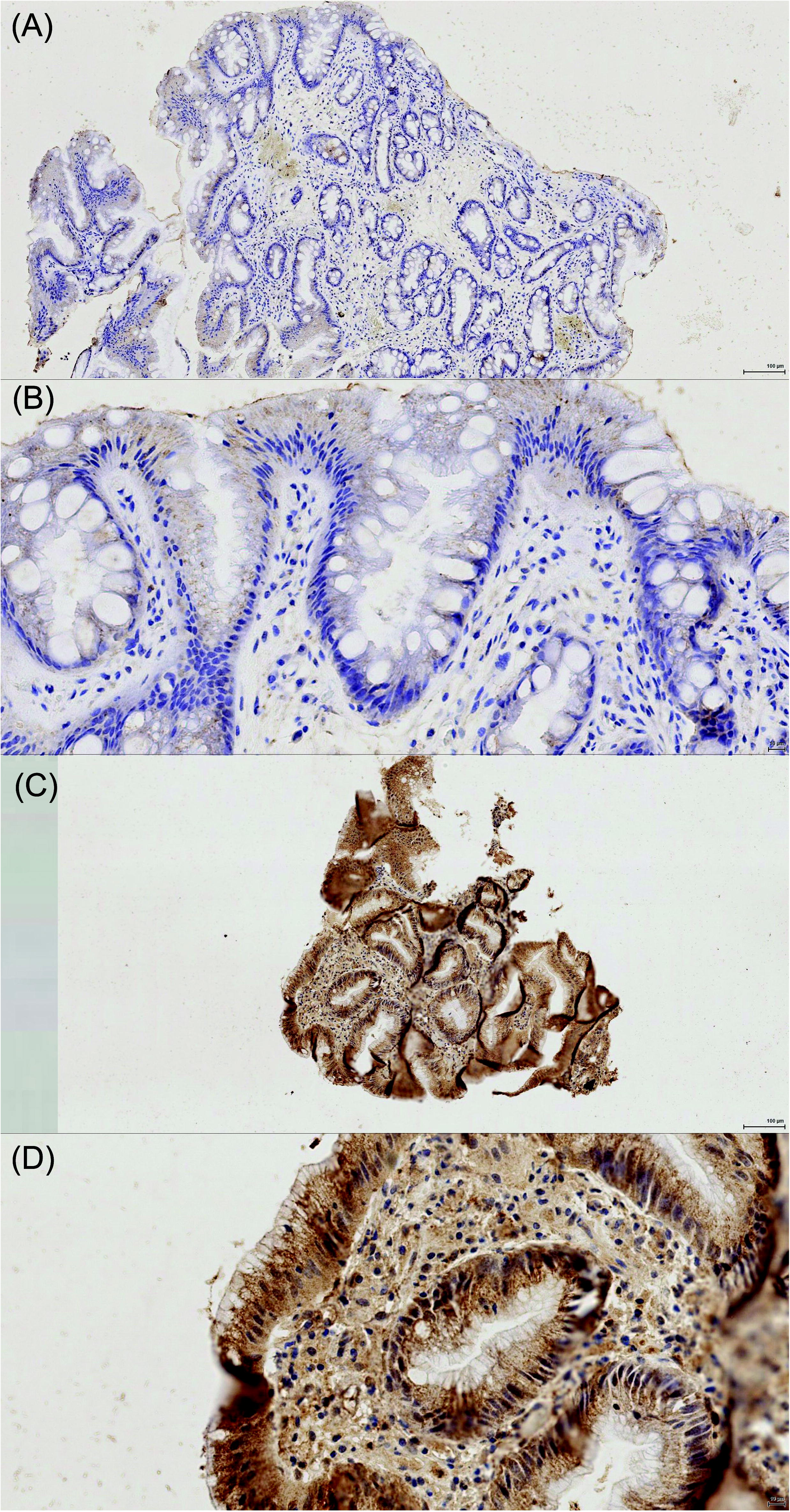

We then validated cg17337672 within FGFR2, finding that median methylation was 83.4% in progressors vs. 82.2 in non-progressors (p=0.51,z=0.653, Wilcoxon rank sum test).

### Expression of OR3A4 in progressors vs. non-progressors

For the stromal compartment, a median expression of 6 (IQR 4-7) was seen in progressors and 2 (IQR 1-3) in non-progressors (Wilcoxon rank sum p=0.0308, z=−2.160). For the epithelial compartment, a median expression of 8 (IQR 8-10) was seen in progressors and 5.5 (IQR 3-10) in non-progressors (Wilcoxon rank sum p=0.4587, z=−0.741). Percentage methylation at OR3A4 and stromal expression was strongly negatively correlated (Pearson correlation coefficient = -0.85, p=0.014) but with epithelial expression correlated negatively but non-significantly with methylation at OR3A4 (Pearson correlation coeffiecient = -0.40, p=0.373).

### Ability of OR3A4 methylation to act as a discriminator in BO

In order to understand the accuracy of using methylation within cg09890332 of OR3A4 as a biomarker for high risk Barretts oesophagus we further carried out a multivariate reverse stepwise logistic regression analysis of methylation at the three tagged CpG dinucleotides within the pyrosequencing assay as the independent variables and progressor vs. non-progressor status status as the dependent variable. In this model, CpGs two and three became non-significant (p=0.3325 and p=0.4764) and were removed from the model, leaving the first CpG in cg09890332 as being significant (coef =−0.0563, SE=0.016, z=−3.40,p=0.001,95% CI -0.089- -0.024). Using ROC modelling the AUC of this model was 0.82 (95% CI 0.80-0.83).

We then further used the *<u>diagt</u>* function of Stata 11.2 to model a set methylation threshold effect on sensitivity and specificity of the test, aiming for maximum negative predictive value and correcting for a incidence rate within the cohort of 0.7%. Modelling at a threshold of below 89% being significant showed that hypomethylation at OR3A4 can predict progression to invasive carcinoma with a sensitivity of 70.8%, specificity of 86%, positive predictive value of 85% and negative predictive value of 72.5%.

## CONCLUSIONS

We have identified that hypomethylation at cg09890332 corresponding to the CG nucleotide at position (CHR) of *OR3A4* can discriminate patients who progress from non-dysplastic Barrett’s oesophagus to those who did not. This association is maintained across independent cohorts, and seems to be related temporally as well as by case status. We observed hypomethylation within these samples up to 13 years before the development of invasive adenocarcinoma suggesting this is an early event in the development of malignant change.

The effect of hypomethylation on the *OR3A4* gene seems to be functional, in that immunohistochemistry reveals a change in increased *OR3A4* expression in samples with hypomethylation. The finding that the stromal expression in particular is increased is of interest given the known effect of “pathological” stroma in the pathogenesis of oesophageal cancer. Our observed region co-incides within 225bp of a CTCF and RAD21 transcription factor binding site, further suggesting that methylation there has a function effect to prevent transcription factor binding and alter gene expression. Guo et al [33] performed a genomewide screen of long noncoding RNAs in gastric adenocarcinoma, finding that *OR3A4* was significantly (55.9 fold,) over-expressed in these patients. They also observed that levels of *OR3A4* were correlated with metastatic potential and prognosis. Furthermore, they utilised *OR3A4* over-expression vectors and performed siRNA knockdown to demonstrate that *OR3A4* seems to regulate cellular proliferation in gastric cancer cell lines. Finally, they utilised their over-expressing cell line models and implanted them into nude mice, finding that *OR3A4* over-expressing gastric cancer cell lines grew significantly faster and more aggressively than with knockdown of *OR3A4.* Downstream analysis of target genes demonstrated that *OR3A4* targets *PDLIM2,* a putative tumour suppressor than regulates cell cycle and adhesion; *PIWIL1,* a transcriptional silencer and *DLX4* which induces epithelial-mesenchymal transition via *TWIST1.* Clearly, our observation that OR3A4 is over-expressed via hypomethylation is clearly relevant in the pathogenesis of Barrett’s oesophagus that progresses to invasive adenocarcinoma.

We found both at the individual probe level and as part of a differentially methylated region that there is hypomethylation in the CpG island associated with *FGFR2,* however this did not validate at the single probe level when examined with bisulphite pyrosequencing. *FGFR2* has been observed to undergo recurrent alteration in both oesophageal adenocarcinoma [34] and squamous cell carcinoma [35] with the latter demonstrating recurrent amplification. The disparity between our microarray results and validation by pyrosequencing may be due to probe inflation caused by small sample size, however given its biological associations with oesophageal adenocarcinoma further work is needed.

In common with pre-malignant lesions in cancer, such as colorectal adenomatous polyps [28], we observed a trend towards genome wide hypomethylation suggesting a widespread over-expression of genes as part of the development towards malignancy. We also found no difference in chromosomal instability between progressors and non-progressors, although there was widespread instability within both sets of samples, in common with what has previously been observed [36] in Barretts oesophagus.

This methylation marker, although seemingly highly accurate for the detection of progression of Barrett’s to invasive adenocarcinoma, is likely to be of more utility as a multi-modal stratifier in Barrett’s oesophagus, taking account of previous findings at the mutational and copy number level, as well as epigenetic change. However, for the purpose of designing a surveillance programme with the ability to risk stratify the non-dysplastic Barretts oesophagus patient, this marker has significant potential utility.

Development of a streamlined surveillance programme could lead to cost savings through the avoidance of un-necessary upper gastro-intestinal endoscopy in a low risk cohort identified by molecular marker panel. More frequent endoscopy in the high risk cohort could lead to earlier diagnosis of OADC or initiate management of BO to arrest further progression. In conclusion, development of a stratified marker panel in the context of a clinical trial is now needed to improve diagnosis of high risk BO.

## Contributorship statement

Cohort identification and follow-up: NT, MTH, OT

Molecular analysis: MPD, TN, JDS, CW, LT, JDJ, ADB

Histological processing and analysis: TN, MPD, RH

Manuscript: ADB, TN, MD.

Guarantor: ADB.

